# High fat diet exacerbates cognitive decline in mouse models of Alzheimer’s disease and mixed dementia in a sex-dependent manner

**DOI:** 10.1101/2021.10.05.463111

**Authors:** Olivia J. Gannon, Lisa S. Robison, Abigail E. Salinero, Charly Abi-Ghanem, Febronia Mansour, Alvira Tyagi, Rebekah Brawley, Jordan Ogg, Kristen L. Zuloaga

## Abstract

Approximately 70% of Alzheimer’s disease (AD) patients have co-morbid vascular contributions to cognitive impairment and dementia (VCID); this highly prevalent overlap of dementia subtypes is known as mixed dementia (MxD). AD is more prevalent in women, while VCID is slightly more prevalent in men. Sex differences in risk factors may contribute to sex differences in dementia subtypes. Unlike metabolically healthy women, diabetic women are more likely to develop VCID than diabetic men. Prediabetes is 3x more prevalent than diabetes and is linked to earlier onset of dementia in women, but not men. How prediabetes influences underlying pathology and cognitive outcomes across different dementia subtypes is unknown. To fill this gap in knowledge, we investigated the impact of diet-induced prediabetes and biological sex on cognitive function and neuropathology in mouse models of AD and MxD. Male and female 3xTg-AD mice received a sham (AD model) or unilateral common carotid artery occlusion surgery to induce chronic cerebral hypoperfusion (MxD model). Mice were fed a control or high fat (HF; 60% fat) diet for 3 months prior to behavior assessment. In both sexes, HF diet elicited a prediabetic phenotype (impaired glucose tolerance) and weight gain. In females, but not males, metabolic consequences of a HF diet were more severe in AD or MxD mice compared to WT. In both sexes, HF-fed AD or MxD mice displayed deficits in spatial memory in the Morris water maze (MWM). In females, but not males, HF-fed AD and MxD mice also displayed impaired spatial learning in the MWM. In females, but not males, AD or MxD caused deficits in activities of daily living, regardless of diet. Astrogliosis was more severe in AD and MxD females compared to males. Further, HF diet caused greater accumulation of amyloid beta in MxD females compared to MxD males. In females, but not males, more severe glucose intolerance (prediabetes) was correlated with increased hippocampal microgliosis. In conclusion, high fat diet had a wider array of metabolic, cognitive, and neuropathological consequences in AD and MxD females compared to males. These findings shed light on potential underlying mechanisms by which prediabetes may lead to earlier dementia onset in women.

**Highlights:**

- Created a mouse model of mixed dementia (MxD) with both AD + VCID pathology.
- HF diet caused greater metabolic impairment in AD and MxD females, compared to males.
- AD and MxD females showed a wider array of cognitive deficits, compared to males.
- Astrogliosis and Aβ pathology were more severe in AD/MxD females, compared to males.
- Metabolic impairment was more consistently associated with reductions in cognitive function in females.
- More severe glucose intolerance was associated with worse microgliosis in females only.

## Introduction

Diabetes increases the risk of developing dementia by 2-fold (1–5). While diabetes is increasingly common, prediabetes is estimated to effect 1 out of every 3 Americans (6), and most people are unaware of their status. Like diabetes, prediabetes is characterized by impaired glucose tolerance; however, those with prediabetes show slight elevations in insulin and fasting blood glucose, rather than hyperglycemia. Diabetes and prediabetes are shared risk factors and common co-morbidities for the two most common forms of dementia: Alzheimer’s disease (AD) and vascular contributions to cognitive impairment and dementia (VCID) (1–5). Obesity, which is often comorbid with prediabetes or type 2 diabetes, increases AD risk 3-fold and VCID 5-fold (7–9). AD is characterized by amyloid plaques, tau tangles, and neurodegeneration culminating in brain atrophy and cognitive impairment. VCID is caused by deficits in cerebral blood flow and/or damage to cerebral vessels. In reality, the distinction between AD and VCID is less clear-cut. Dementia pathologies often overlap, with more than half of dementia patients having multiple pathologies (10), a condition known as mixed dementia (MxD). The most common form of MxD is a mix of AD and VCID, as this occurs in ~70% of AD patients (11–13). MxD is underrepresented in animal research despite its high clinical prevalence. Understanding the interaction between AD and VCID risk factors will provide insight into MxD.

Dementia risk and prevalence vary by sex: women are more likely to develop AD (10), while men are slightly more likely to develop VCID (14, 15). Discrepancies in risk factors may drive these sex differences. In patients with diabetes, the sex difference seen in the non-diabetic population is reversed: diabetic women are at a 19% greater risk of VCID than diabetic men (3). Among those who have VCID, women are also more likely to have diabetes (16). Prediabetes is associated with cognitive impairment and earlier onset of dementia in women, but not men, suggesting it may be a sex-specific risk factor (17). However, it is unknown how prediabetes affects MxD, and whether these effects differ by sex.

High fat (HF) diet is commonly used to induce metabolic disease in rodents, as it causes both obesity and prediabetes. HF diet can have profound effects on the brain, some of which are sex-dependent. For example, we have shown that HF diet impairs adult hippocampal neurogenesis in female, but not male mice (18). Further, we recently found that HF diet in middle aged mice causes a wider array of cognitive deficits in females compared to males (19). Others have found that a HF diet in the 3xTg-AD model of AD exacerbates cognitive impairment (20–25) and AD pathology such as brain atrophy (20), inflammation (22, 26), and Aβ load (26–28). Examination of sex differences in the cognitive effects of HF diet in the 3xTg-AD mouse have been mixed, with some studies finding greater cognitive impairment in females (20), while others have found no sex differences (21). Previously, we reported that HF diet results in greater metabolic impairment (weight gain, visceral fat, and glucose intolerance) in female compared to male 3xTg-AD mice (29). HF-fed females also showed increased astrogliosis in the hypothalamus, a brain region that controls metabolic function. Whether these greater metabolic disturbances in females would also lead to more severe cognitive deficits and neuropathology in brain regions associated with learning and memory had not yet been tested. In the current study, using mouse models of both AD and MxD, we found that diet-induced obesity with prediabetes led to a wider array of cognitive deficits and neuropathology in females compared to males.

## Methods

### Animals and Experimental Design

This study was conducted in accordance with the National Institutes of Health Guidelines for the Care and Use of Laboratory Animals, and protocols were approved by the Institutional Animal Care and Use Committee at Albany Medical College (Albany, NY, USA). Temperature and humidity were set at 72 °F, 30-70% humidity, with a 12 h light/dark cycle (7 a.m. on/7 p.m. off). Mice were fed a standard chow diet (Purina Lab Diet 5P76) until the start of the study. They were housed in Allentown cages at a density of 2-5 mice. Mice were provided with environmental enrichment (Nestlets and Shepherd Shacks) and were group housed at all times, except during the nest building test. Male and female wild-type (WT) B6129SF2/J mice (#101045) and 3xTg-AD (#34830-JAX) breeding pairs were purchased from Jackson Laboratories (Bar Harbor, ME) and used to maintain a colony at Albany Medical Center’s Animal Resource Facility. The 3xTg-AD mice, which are on a C57BL/6;129X1/SvJ;129S1/Sv background, have three mutations that are associated with AD in humans: APPSwe, tauP301L, and Psen1^tm1Mpm^ (30). A timeline of the experiment is shown in **Figure 1A**. At ~3 months of age, 3xTg-AD mice underwent a sham surgery (AD group) or a unilateral common carotid artery occlusion surgery (MxD group). WT controls also received a sham surgery (WT group). One week following surgery, mice were placed on either a HF diet (60% fat, 5.24 kcal/g; D12492, Research Diets, New Brunswick, NJ) or a low-fat (LF) control diet (10% fat, 3.82 kcal/g; D12450B, Research Diets) for the duration of the study. Approximately 2.75 months later, mice underwent a glucose tolerance test (GTT), a 2-week rest period, behavioral testing, blood flow imaging, euthanasia, and tissue collection (including brains, fat, and reproductive organs). A total of 251 mice were used in this study. Experiments were conducted in cohorts of up to 20 animals. A subset of mice (n=118) was designated for behavior testing. In total, 23 mice were excluded due to premature death or the presence of other major health exclusions (hydrocephaly, large fighting wounds, tumors). Final group sizes ranged from 13-25 per group for metabolic measures, 8-13 for behavioral testing, and 4-6 for immunohistochemistry (IHC). During tests, experimenters were blinded to surgical group. Blinding to diet and sex were not possible due to mouse appearance. During analysis, experimenters were blinded to sex, diet, and dementia group.

**Figure 1.**
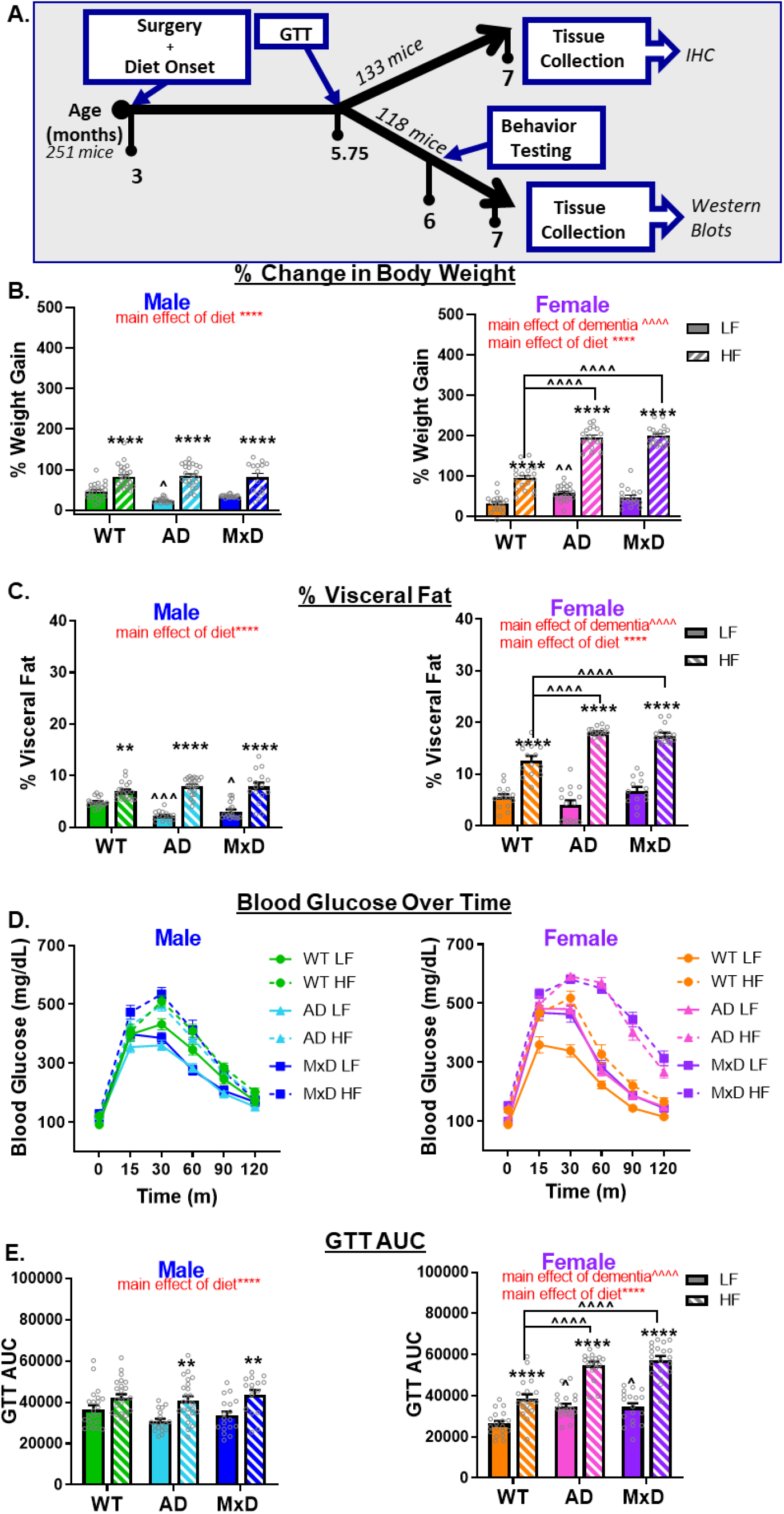
HF diet caused greater metabolic impairment in AD and MxD females compared to males. A) Experimental timeline. GTT (glucose tolerance test). B) Weight gain was assessed by the % change in body weight from the start of the study to the end of the study. C) Visceral adiposity was determined by isolating and weighing the visceral fat pads and normalizing to body weight. D, E) Glucose intolerance was assessed with a GTT following a 16hr fast. D) Glucose clearance was gauged by concentrations of glucose in the blood measured over time (time 0= fasting blood glucose). E) Blood glucose concentration over time was used to calculate area under the curve. We previously reported metabolic data for the Sham WT and Sham AD, but not the MxD, groups in Robison et al (2020) in the *Journal of Neuroinflammation* (29); licensed under a Creative Commons Attribution 4.0 International License; https://creativecommons.org/licenses/by/4.0/). **p<0.01 effect of diet, ****p<0.0001 effect of diet, ^p<0.05 effect of dementia, ^^^p<0.001 effect of dementia, ^^^^p<0.0001 effect of dementia. Data are presented as mean + SEM (n=13-25/group).

### Surgical model of MxD

To model MxD, 3xTg-AD mice underwent a right common carotid artery occlusion surgery, as previously described, to elicit chronic cerebral hypoperfusion and model VCID (Zuloaga, 2016; Zuloaga, 2015; Salinero 2020). Briefly, under isoflurane anesthesia, the right common carotid artery was ligated with two 6-0 silk sutures and cauterized (MxD group). The sham surgery (WT and AD groups only) consisted of exposing the carotid artery without ligation. Incision sites were closed with Vetbond, and the mice were given 100μL 0.03mg/mL buprenorphine via subcutaneous injection twice per day for 3 days as an analgesic.

### Glucose Tolerance Test

As previously described (19, 29, 31), mice were given a glucose tolerance test (GTT) to assess diabetic status after being on either the control or HF diet for 3 months. The mice were fasted overnight, and their fasting blood glucose levels were measured (t=0) using a glucometer (Verio IQ, OneTouch, Sunnyvale CA, USA) from their tail vein. Following an i.p. injection of 2g/kg of glucose, blood glucose levels were measured at 15, 30, 60, 90, and 120 min post-injection to assess glucose tolerance.

### Behavior Testing

Following a two-week recovery post-GTT, mice were tested for exploratory activity and anxiety-like behavior in the open field (day 1), episodic-like memory in the novel object recognition test (NORT; day 2), spatial learning and memory in the Morris water maze (MWM; days 8-10 or 10-12), and activities of daily living using a nest-building task (days 15-16). Videos were recorded of behavioral performance for open field, NORT, and MWM and analyzed using automated tracking software (ANY-maze 5.1, Stoelting, Wood Dale, IL). For each test, mice were placed into the procedure room under dim light and allowed to acclimate for 1 hour. Each test apparatus was cleaned with 70% ethanol between each mouse to remove olfactory cues.

#### Open Field

The mice were placed in the test apparatus (495 x 495mm box) for 10 minutes. Distance traveled was used to determine the general activity levels of the animal. The percent of time spent in center of the arena was used to determine anxiety-like behavior.

#### NORT

NORT consisted of two, five-minute trials performed in the same open field arena. In the first trial, mice were placed in the box and allowed to explore two identical objects (rubber ducks). Mice were then returned to a recovery cage for 1 hour. For the second trial, mice were returned to the arena, with the one familiar object replaced with a novel object (salt shaker). Episodic-like memory was assessed by recognition index [(time with novel object/total time with objects)*100]. Between tests, objects were cleaned with 70% ethanol to mask olfactory cues. Mice that spent less than ≤2 seconds with the objects were excluded.

#### MWM

Hippocampus-dependent spatial learning and memory were assessed using a modified 3-day version of the MWM that has been shown to be optimal for older, cognitively impaired, obese mice (32). The protocol has been previously described in detail (19). On day 1, 5 visible trials were performed in which mice learn to find the platform with a visual cue (flag). The entry point was alternated for each trial. On day 2, mice underwent 5 hidden trials, in which the visual cue was removed from the platform. All trials were 3 minutes long with a 30-minute intertrial interval. The distance traveled to reach the platform (pathlength) was used as a measurement of non-spatial (visual trials) and spatial (hidden trials) learning. On day 3, a single probe trial was performed in which the platform was removed from the pool. Spatial memory was calculated as the percent of time spent in the target quadrant of the pool during the first minute of the probe trial.

#### Nest Building

Mice were singly housed in Allentown cages with pine chip bedding and two pre-weighed Nestlets each. After 16 hours (overnight), the mice were removed from their test cage and returned to group housing. Nests were rated on a 1-5 scale (with half-point scores allowed) based on published criteria (Deacon, 2006) by 3 experimenters that were blinded to treatment group. The 3 ratings were averaged.

### Cerebral Blood Flow Measurement

Cerebral blood flow was measured via laser speckle contrast imaging (moor FLPI full field laser perfusion imager; Moor Instruments, Wilmington, DE, USA) under isoflurane anesthesia as previously described (19). Image acquisition (5-minute scan), processing, and analysis were performed using moorFLPI Review V4.0 software (Moor Instruments, Wilmington, DE, USA). Average flux values were extracted from regions of interest (ROIs) using a published protocol (Poloycarpou, 2016). Measurements are presented as %difference in blood flow between the left (non-ischemic for MxD mice) and right (ischemic for MxD) hemisphere.

### Immunofluorescence

Mice were perfused with ice-cold 0.9% saline. Brains were removed and fixed in 4% paraformaldehyde for 24 hours, followed by immersion in 30% sucrose for at least 72 hours. Brains were then snap frozen in OCT and stored at −80°C until sectioning. Brains were sectioned at 40-microns on a Leica CM1950 cryostat into 6 series. Sections were washed in PBS containing 0.01% sodium azide, permeabilized at room temperature for 1 hour (0.3% TPBS) and blocked for 1 hour at room temperature in 4% donkey serum in 0.3% TPBS before being incubated in blocking buffer with primary antibodies at 4 °C overnight. Primary antibodies for one series included rabbit anti-beta Amyloid (1:300, Cat# 715800, Lot# SH257822; Invitrogen, Waltham, MA). Primary antibodies for another series included rat anti-glial fibrillary acidic protein (1:2500, AB5804, Millipore, Lot # TA265137) or goat anti-Iba1 (1:1000, PA5-18039, Lot #TI2638761, SJ2467805; ThermoFisher, Waltham, MA). Sections were incubated with secondary antibodies and DAPI (1:1000, Cat# D1306, ThermoFisher, Waltham, MA) in blocking buffer for 2 hours at room temperature. Secondary antibodies used included Rhodamine Red-X Donkey Anti-Rabbit (1:100), Alexa Fluor 647 Donkey Anti-Goat (1:300), DyLight™ 405 AffiniPure Donkey Anti-Rat (1:300) (Jackson ImmunoResearch, West Grove, PA), Alexa Fluor® 647 AffiniPure Donkey Anti-Goat IgG (H+L) (1:1000, Cat# AB_2340437, Jackson ImmunoResearch, West Grove, PA), and Alexa Fluor® 488 Donkey Anti-Rabbit (1:1000, Cat# ab150073, Jackson ImmunoResearch, West Grove, PA). Images for quantification were taken at 10x using the Axio Observer fluorescent microscope (Carl Zeiss Microscopy, Jena, Germany). All analyses were performed in the right hemisphere of coronal sections by an experimenter who was blinded to treatment group. For Aβ quantification, regions of interest (ROI) were drawn in the cortex and number of cells positive for Aβ were counted **(Figure 4C**). For neuroinflammation analysis (microglia and astrocytes), ROIs were drawn in CA1, CA2, CA3, and dentate gyrus of the hippocampus in ~3 brain sections per mouse and % area covered was measured using ImageJ.

**Figure 4.**
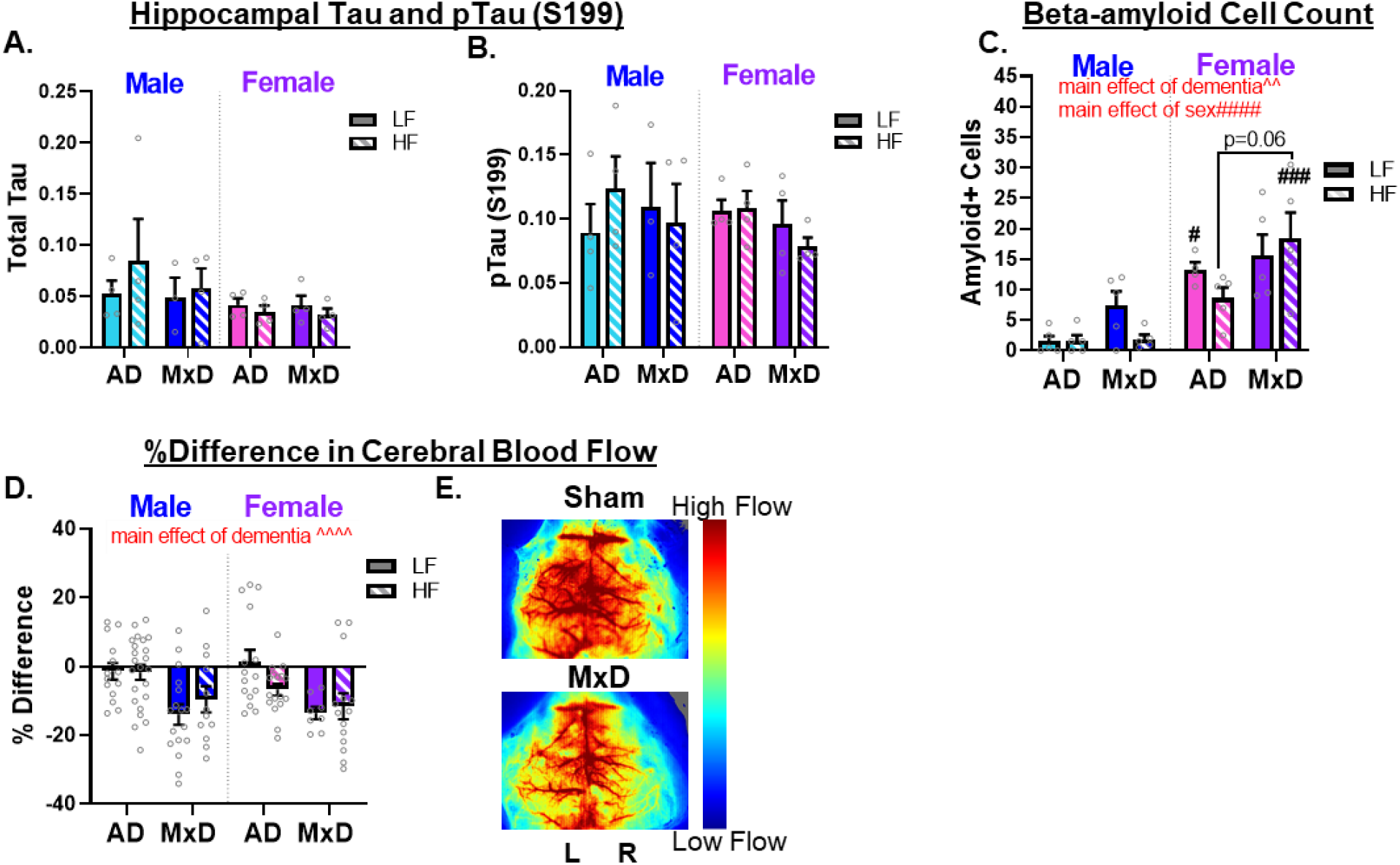
MxD mice exhibit deficits in blood flow regardless of sex, and female AD/MxD mice have greater Aβ, but not tau, pathology. Western blot was used to determine the amount of tau protein in right (ischemic for MxD) hippocampal isolates. Measurements for A) total tau, and B) pTau S199 were normalized to GAPDH. Data are presents as mean + SEM (n=3-4/group). C) To examine amyloid burden, the number of cells positive for Aβ in a cortical region of interest were counted. Data are presents as mean + SEM (2-way ANOVA with Tukey’s multiple comparison test, effect of sex # p<0.05, ### p<0.001, n=4-5/group). D) Cerebral blood flow was measured in the temporal region using laser speckle blood flow imaging 4 months following unilateral common carotid artery surgery (MxD). The % difference in blood flow between the ischemic and non-ischemic hemispheres with a value closer to 0 indicating no difference in blood flow and a negative % difference indicating lower blood flow in the hemisphere ipsilateral to the occlusion. E) Representative images for blood flow scans show cerebral blood flow for sham (top) and MxD (bottom). Data are presents as mean + SEM (^^^^ main effect of dementia p<0.0001, 3-way ANOVA, n=8-23/group, % difference in temporal blood flow).

### Western Blot

Frozen hippocampi (ipsilateral to VCID or sham surgery) were thawed and homogenized in 50μL RNA-Later (45-R0901-100MLsigma). 25μL of that homogenate was transferred into 100 μL T-Per buffer (ThermoScientific #78510) supplemented with protease and phosphatase inhibitor cocktail (HALT, ThermoScientific #1861284) and spun at 21000g for 20 minutes. The supernatant was collected, and the protein concentration was determined using a Pierce BCA Protein Assay Kit (Thermo Scientific #23227). 20μg of protein were denatured in LDS (4X Bolt™ LDS Sample Buffer, Invitrogen, B0007) with 0.2M DTT and boiled for 5 minutes at 95°C. Proteins were then separated using a 10% Tris Bis gel (ThermoFischer, NW00107BOX) before being transferred onto a nitrocellulose membrane (Nitrocellulose/Filter Paper Sandwich, 0.2 μm, 8.3 x 7.3 cm, ThermoFischer, LC2000). Membranes were blocked using LI-COR blocking solution (Intercept® (TBS) Blocking Buffer, LI-COR, 927-60001) for one hour at room temperature then exposed to primary antibodies at 4°C overnight. The following day membranes were washed then incubated with the corresponding secondaries before being washed and scanned using an Odyssey CLx LI-COR scanner. All membranes were scanned simultaneously to ensure similar exposure and scanning parameters. Membranes were then stripped using LI-COR stripping buffer (NewBlot™ IR Stripping Buffers for NIR Western Blots, LI-COR, 928-40028) for 20min and rescanned to ensure loss of signal before being incubated with an anti-GAFPH antibody (Sigma, 45-G9545-100UL) as a loading control. Western blot band analysis was conducted using the LI-COR Image Studio and normalized to loading control. Antibodies used were for total tau ((T46) mouse abCam203179, lot #GR33658561) and phosphorylated tau (S199-202, rabbit Invitrogen 44-768G lot SG255287). All primary antibodies were used at 1:1000 in Tris buffered saline supplemented with 0.1% Tween (TBST). Secondary antibodies were used at 1:10,000 in TBST with 0.02% SDS. Secondary antibodies included IRDye® 800CW Donkey anti-Mouse IgG (CAT# 926-32212, Lot#D00930-09, LI-COR, Lincoln, Nebraska) and IRDye® 680RD Donkey anti-Rabbit IgG (CAT# 926-68073, Lot#D00421-09, LI-COR, Lincoln, Nebraska) were used at 1/10000 in TBST with 0.02% SDS.

### Statistics

Statistical analyses were performed using Prism 8.1 (GraphPad Software, San Diego, CA, USA). All data is presented as mean +SEM, with significance set at p<0.05 except for categorical data (nest building is expressed as median + interquartile range). Following a Grubbs’ test for statistical outliers, a 2-way ANOVA was performed with Tukey’s correction for multiple comparisons [dementia type (WT vs. AD vs. MxD) X diet (LF vs. HF)] in data segregated by sex. In secondary analyses to assess sex differences, 3-way ANOVAs were performed (sex X dementia type X diet). In measures with large sample sizes (n ≥13 for all groups) a ROUT test was performed (metabolic data only). One-sample t-tests were performed for measurements where values are compared to chance (50% in NORT, 25% in MWM probe trial, and 0% hemispheric difference in blood flow).

## Results

### Animal models and timeline

In order to create a mouse model of MxD, 3xTg-AD mice (~3 months of age) underwent a unilateral common carotid artery occlusion surgery to induce chronic cerebral hypoperfusion/vascular pathology. These mice were compared to AD mice (3xTg-AD that underwent a sham surgery) and WT controls (WT mice that underwent a sham surgery). In order to determine the effects of HF diet-induced prediabetes on outcomes in the AD and MxD models, one week following surgery, mice were placed on either a HF diet (60% fat) or a control LF diet (10% fat) from ~3-7 months of age. A study timeline is shown in **Figure 1A**.

#### HF diet caused greater metabolic impairment in AD and MxD females compared to males

We have previously reported (29) that metabolic effects of a HF diet are more severe in 3xTg-AD females compared to males. Here we show that these sex differences persist in the MxD model. Within each sex, there was a main effect of HF diet to increase % weight gain (p<0.0001; **Figure 1B**), visceral fat accumulation (p<0.0001; **Figure 1C**), and glucose intolerance (p<0.0001; **Figure 1D, E**). Dementia X diet interactions were sex-dependent. In females, post-hoc tests showed that HF-fed AD and HF-fed MxD females had greater metabolic impairment (weight gain, visceral fat accumulation, and glucose intolerance) than HF-fed WT females (p<0.0001 for all measures). Conversely, in males, LF-fed AD and LF-fed MxD males had less visceral fat than LF-fed WT males (p<0.001 for AD, p<0.05 for MxD). Analysis of sex differences via a 3-way ANOVA (**Supplemental Table 1**) showed a sex X dementia interaction and sex X diet interaction (p<0.0001 for each interaction) in which both visceral fat and glucose intolerance were exacerbated in AD/MxD and HF-fed females, but not males. Further, there was also a sex X diet X dementia interaction for % weight gain (p<0.0001), which was driven by the large % weight increase in HF-fed AD and HF-fed MxD females. Taken together, the data show that HF diet caused greater metabolic impairment in AD or MxD females compared to WT females or AD or MxD males.

### HF diet caused a wider array of cognitive deficits in females

Our prior work has demonstrated that HF diet causes a wider array of cognitive deficits in middle aged females, compared to males, in a mouse model of VCID (19). Whether HF diet would differentially impact AD and MxD in each sex was unknown. Preference for the novel object in the NORT (episodic-like memory; **Figure 2A**) was assessed with a one sample t-test vs. no preference (recognition index of 50%). In males, all WT mice demonstrated a preference for the novel object (LF-fed WT p<0.001, HF-fed WT p<0.001). In females, LF-fed WT mice show intact memory (p<0.001), while HF-fed WT females only showed a trend towards preference for the novel object (p=0.09). In both sexes, both AD and MxD groups on either diet demonstrated no preference for the novel object, indicating an impairment in episodic-like memory. A 3-way ANOVA showed no sex differences. Although there were some group differences in exploratory behavior and anxiety-like behavior in the open field, neither of these measures correlated with NORT performance (**Supplemental Figure 1C**). The MWM was used to examine spatial learning and memory. All groups were trained to swim to a visible platform (main effect of trial p<0.05, **Supplemental Figure 1E**). When the platform was hidden (spatial learning trials; **Figure 2B, C**), no group differences were observed in males. In females, there was a dementia x diet interaction (p<0.05), in which HF-fed AD females, LF-fed MxD females, and HF-fed MxD females all had significantly impaired spatial memory compared to LF-fed WT females (p<0.01 for each group; post-hoc tests). A 3-way ANOVA (**Supplemental Table 1**) showed a sex X dementia X diet interaction (p<0.01) in which HF-fed AD/MxD females had the most severe spatial learning deficits. In the probe trial (spatial memory; **Figure 2D**), preference for the target quadrant was assessed with a one-sample t-test of % time in target quadrant vs. chance (25%). In males, only WT mice (on either diet) and LF-fed AD mice showed a preference for the target quadrant (p<0.05), indicating that spatial memory was impaired in HF-fed AD males and MxD males on either diet. In females, only the LF-fed WT mice showed a preference for the target quadrant (p<0.05), indicating that HF-fed WT females and AD or MxD females on either diet had impaired spatial memory. A 3-way ANOVA (**Supplemental Table 1**) showed only a slight trend towards worse spatial memory in females (p=0.08) with no significant interactions. Although there were some group differences in swim speed, this measure did not correlate with learning or memory in the MWM (**Supplemental Figure 1D**). The nest building test (**Figure 2E**) was used to assess activities of daily living (ADLs). In males, all mice performed well although there was a main effect of diet (p<0.05) to impair ADLs. In females, AD and MxD females showed clear impairments (main effect of dementia p<0.0001), with LF-fed MxD (p<0.01) and HF-fed MxD (p<0.05) females building poorer nests than control LF-fed WT females. HF-fed AD females also showed a trend (p=0.056) towards impaired ADLs. A 3-way ANOVA showed a main effect of sex X dementia interaction (p<0.01) in which AD and MxD females were more impaired than males. Taken together, these data show a wider array of cognitive deficits in AD or MxD females compared to males, particularly when females were fed a HF-diet.

**Figure 2.**
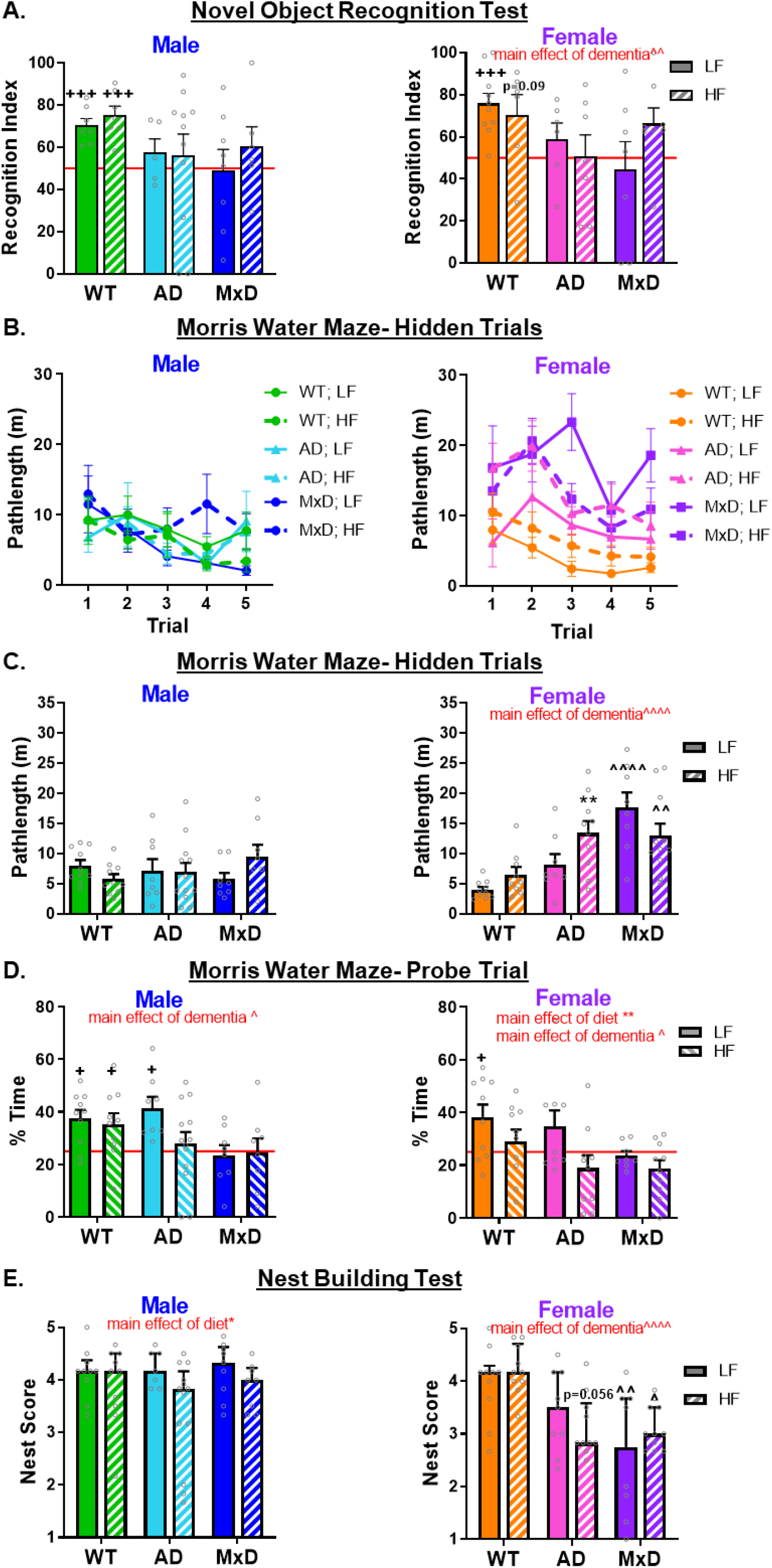
HF diet caused a wider array of cognitive impairment in females compared to males. A) Episodic-like memory was assessed in the novel object recognition test (NORT). Recognition index (% time spent with the novel object) was calculated. Performance not significantly greater than chance (50%, indicated by the red line) indicates impaired memory. B-D) Spatial learning and memory was assessed using the Morris Water Maze (MWM). Five hidden trials (B) assessed spatial learning via pathlength to reach the target platform (shorter pathlength = better performance). Average pathlength over the 5 hidden trials (C) was longer (more impaired memory) in MxD females and AD females on a HF diet. Spatial memory was assessed in the probe trial (D), as % time spent in the target quadrant vs. chance (25%, indicated by the red line). Performance above 25% indicates intact memory. E) The nest building task was used to assess activities of daily living. Nests were graded on 1-5 scale (average of scores by 3 experimenters blinded to treatment). Lower scores are indicative of impairment. +++p<0.001 vs chance, +p<0.05 vs. chance, **p<0.01 effect of diet, ^p<0.05 effect of dementia, ^^p<0.01 effect of dementia, ^^^^p<0.0001 effect of dementia, Red line =chance. Data are presented as mean + SEM, except for nest building (median + interquartile range) (n=5-11/group NOR, n=8-13/group MWM, nest building).

### Neuropathology was exacerbated in AD/MxD females

We have previously reported sex-dependent effects of HF diet on hypothalamic neuroinflammation in 3xTg-AD mice; however, neuroinflammation in brain areas associated with cognition and other forms of neuropathology had not been assessed. Astrogliosis (immunolabeling for glial fibrillary acid protein; GFAP; **Figure 3**) and microgliosis (immunolabeling for Iba-1: **Figure 3**) were assessed in the hippocampal CA1, CA2, CA3 and dentate gyrus regions. In males, astrogliosis (GFAP % area covered) was unaffected by either diet or dementia in any of the regions assessed. In females, there was a main effect of dementia, in which AD and MxD females showed increased hippocampal astrogliosis (CA1 p<0.05, CA2 p<0.01, dentate gyrus p<0.01). A 3-way ANOVA showed a sex X dementia interaction in which AD/MxD females had greater astrogliosis than males (CA1 p<0.05, CA2 p<0.05, CA3 p<0.01, dentate p<0.05). In males, there was a main effect of dementia on microgliosis (Iba1 % area covered), in which AD and MxD males showed decreased hippocampal microgliosis in several regions (CA1 p<0.001, CA2 p<0.0001, CA3 p<0.01). In females, microgliosis was unaffected by either diet or dementia in any of the regions assessed. A 3-way ANOVA showed a sex X dementia interaction in several regions (CA1 p<0.01, CA2 p<0.01, CA3 p<0.05), which was driven by higher microgliosis in WT males (regardless of diet) compared to all other groups. To examine AD pathology, we assessed phosphorylated tau in the hippocampus via Western blot (**Figure 4A, B; Supplemental Figure 2**) and Aβ positive cells in the cortex via immunolabeling (**Figure 4C**). No differences in phosphorylated or total tau were detected between AD and MxD groups, regardless of sex or diet; however, differences were detected in Aβ. In males, there was a main effect of MxD to increase Aβ (p<0.05) that was driven by increased Aβ in LF-fed MxD males. In females, there was a trend for a main effect of MxD to increase Aβ (p=0.068). A 3-way ANOVA showed that AD/MxD females had higher levels of Aβ-positive cells than males (p<0.0001) and that there was a main effect of MxD to increase Aβ (p<0.01). To validate that the unilateral carotid artery occlusion surgery modeled VCID by inducing chronic cerebral hypoperfusion, cortical blood flow was measured using laser speckle contrast imaging 4 months following surgery (**Figure 4D**). As expected, blood flow deficits were found in the right (occluded) hemisphere of all MxD groups (main effect of MxD, p<0.0001). No sex differences were detected. Taken together, this data shows that although some pathology was equivalent between the sexes, hippocampal astrogliosis and cortical Aβ pathology were more severe in AD/MxD females, compared to males.

**Figure 3.**
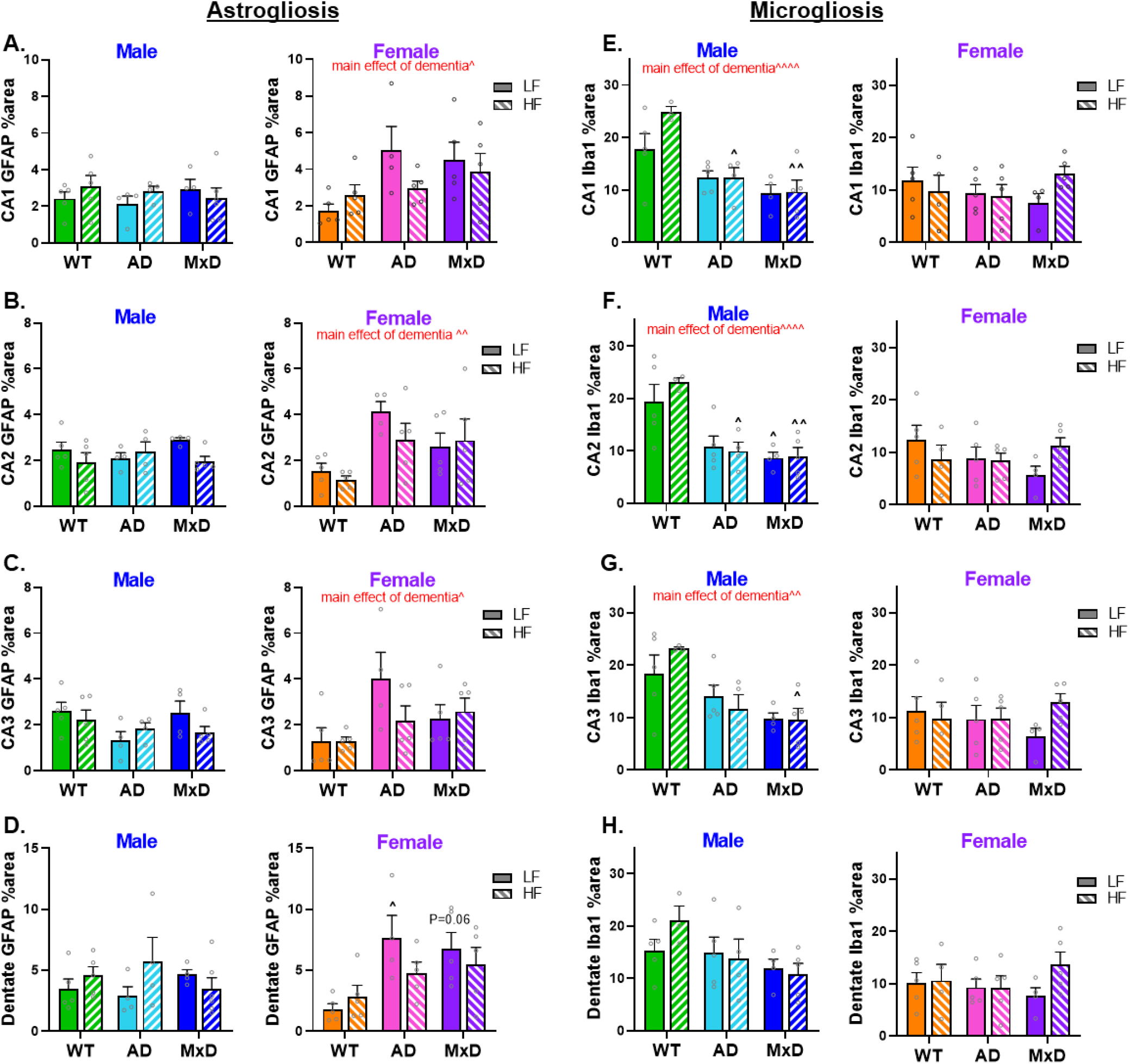
Astrogliosis is exacerbated in AD/MxD females, while microglia coverage is decreased in AD/MxD males. Astrogliosis in multiple regions of the hippocampus was gauged through GFAP immunofluorescence (greater % area covered indicating greater astrogliosis). Hippocampal regions of interest examined: CA1 (A), CA2 (B), CA3 (C), and the dentate gyrus (D). Microglia coverage in the CA1 region of the hippocampus was gauged through Iba1 immunofluorescence (larger % area covered indicating greater microgliosis). Iba1 immunoreactivity was used to calculated microglia coverage as the percent area covered by Iba1. Hippocampal regions of interest examined: CA1 (E), CA2 (F), CA3 (G), and the dentate gyrus (H). Data are presented as mean + SEM (effect of dementia: ^ p<0.05, ^^p<0.01 2-way ANOVA, n=4-5/group).

#### Sex-dependent correlations between cognitive performance and metabolic measures

To further examine the relationship between cognitive, metabolic, and neuropathology measures, we performed linear regression analyses and correlation analyses in males and females separately (**Figure 5**). In both males and females, each metabolic measure was strongly correlated with the others (visceral fat, % weight gain, and AUC; r=0.7-0.9, p<0.0001). In males, the effects of metabolic impairments on cognitive function were mixed. In males, higher % weight gain and more severe glucose intolerance (GTT AUC) were both associated with poorer spatial memory (MWM % time in the target quadrant; p<0.05 for each). Additionally in males, % weight gain was associated with reduced ability to perform activities of daily living (nest score; p<0.05); however, more severe glucose intolerance showed a surprising association with better episodic-like memory (NORT recognition index; p<0.05). In females, metabolic impairment showed a consistent association with worse cognitive function. In females, higher% weight gain or more visceral fat were associated with poorer spatial memory (p<0.01 for weight gain, p<0.05 for visceral fat), and more severe glucose intolerance was associated with reduced ability to perform activities of daily living (p<0.01). In males, the degree of metabolic impairment was not associated with alterations in cerebral blood flow, and blood flow was not associated with cognitive function. In females, metabolic impairment was associated with reduced cerebral blood flow (% blood flow in the right hemisphere; p<0.05 for % weight gain, visceral fat, and glucose intolerance). Additionally in females, greater reduction in cerebral blood flow was associated with more severe deficits in the ability to perform activities of daily living (nest building). Taken together, this data shows that impairment in metabolic measures were more consistently associated with reductions in cognitive function in females.

**Figure 5.**
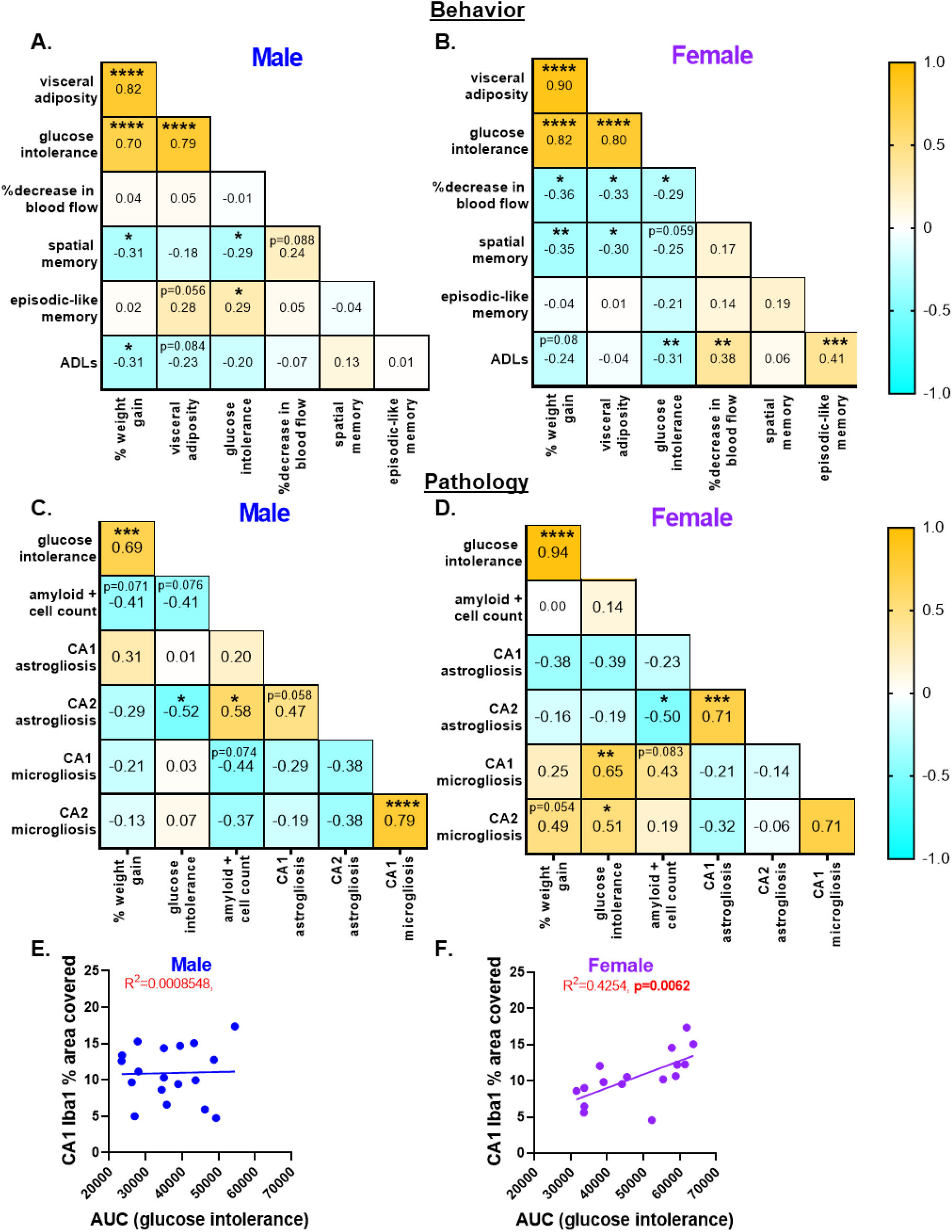
Cognitive impairments and pathological outcomes are correlated with metabolic deficits in male and female mice. Using a correlation matrix, we compared relationships between cognitive, metabolic, and pathological factors. A) Correlations between cognitive and metabolic measures. Visc fat: visceral fat pad weight normalized to body weight (n=55-58/sex); AUC: area under the curve from the glucose tolerance test, high AUC indicates greater glucose intolerance (n=54-58/sex; %decrease in blood flow: the percent difference in cerebral blood flow in the temporal region of the cortical brain surface (n=50-51/sex); MWM % target: % of the time spent in the target quadrant of the probe trial of the MWM test, higher percentage indicates better spatial memory (n=58/sex); NOR% time: % time spent with the novel object in the testing trial of the NOR test, higher percentage indicates better episodic-like memory (n=46-49/sex); ADLs: activities of daily living assessed by score in the nest building test, a lower score indicates worse ADLs (n=57-58/sex). B) Correlations between metabolic and pathological measures in the subset of mice designated for IHC. AUC: area under the curve from the glucose tolerance test, high AUC indicates greater glucose intolerance (n=18-21/sex); amyloid count: the number of cells in a cortical ROI that were positive for beta-amyloid (n=19-20/sex); CA1 GFAP: the % area covered by GFAP staining in the CA1 region of the hippocampus, greater coverage indicates greater astrogliosis in that region (n=18-19/sex); CA2 GFAP, (n=17-19/sex); CA1 Iba2: the % area covered by Iba1 staining in the CA1 region of the hippocampus, greater coverage indicates greater astrogliosis (n=17-18/sex) CA2 Iba1, (n=18/sex). *p<0.05, **p<0.01, ***p<0.001, ****p<0.0001, significant correlation; Pearson r values are presented. Yellow: positive correlation, Blue: negative correlation. C. Linear regression of CA1 Iba1% area covered and glucose tolerance test from the AUC (males: n=19; females n=17). R^2^ value and p value are presented.

**Figure 6.**
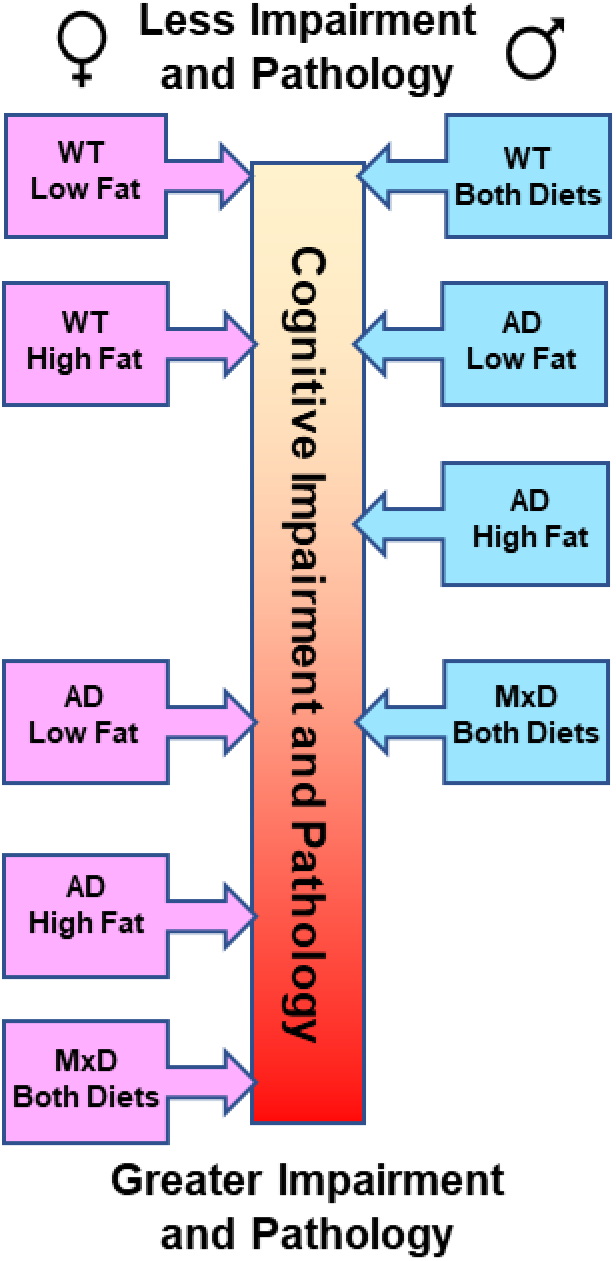
Summary of our major findings. We demonstrate relative accumulation of cognitive impairments and neuropathology of female (left, pink) and male (right, blue) mice along a center scale with the top of the scale indicating less impairment and pathology and the bottom of the scale representing greater impairment and pathology.

Next, correlations in neuropathology were examined (**Figure 5B**). In males, greater cortical Aβ burden was associated with more CA2 astrogliosis (p<0.05), while in females it was associated with less CA2 astrogliosis. In males, more severe glucose intolerance was associated with less CA2 astrogliosis (p<0.02), and there were no significant associations with microgliosis. In females, more severe glucose intolerance was associated with more microgliosis in both CA1 (p<0.01) and CA2 (p<0.05), but not associated with astrogliosis. An example of a sex-specific association is shown in **Figure 5C**. Taken together, this data shows that more severe glucose intolerance was the only metabolic parameter that was associated with worse neuroinflammation; and that this increase in neuroinflammation (microgliosis) occurred in females only.

## Discussion

### Executive summary

The overarching goal of this study was to address gaps in knowledge regarding sex differences in mid-life metabolic risk factors for dementia – obesity and prediabetes. Recent clinical evidence suggests that prediabetes is a risk factor for cognitive decline and dementia in women, but not men (33). How sex and prediabetes interact to influence cognitive function and neuropathology in the two most common forms of dementia - AD and MxD (co-morbid VCID + AD pathology) is unknown. To fill this gap in knowledge, we combined mouse models of VCID (chronic cerebral hypoperfusion) and AD (3xTg-AD mice) to model MxD. We used a chronic HF diet to model obesity and prediabetes. While HF diet ubiquitously caused metabolic impairments, this was augmented in AD/MxD females, but not males. AD, MxD, and HF diet differentially affected cognition in males and females. While episodic-like memory deficits were observed in AD and MxD mice of both sexes, additional cognitive impairments were observed in females. AD/MxD females also showed impairments in activities of daily living, but males did not. When fed a HF diet, AD and MxD females, but not males, also exhibited deficits in spatial learning. Metabolic impairment was also more consistently associated with reductions in cognitive function in females. Moreover, AD and MxD females had more hippocampal astrogliosis and a greater burden of cortical Aβ than males. Finally, more severe glucose intolerance was associated with worse microgliosis in females only. Taken together, the data demonstrate heightened susceptibility of females to the negative metabolic, cognitive, and neuropathological effects of AD, MxD, and HF diet. This data, along with recent clinical findings (33), suggests that prediabetes might contribute to multiple forms of dementia in women.

### Metabolic findings

Here, we found that AD and MxD females had more adverse metabolic responses to chronic HF diet. We have previously shown that sex differences in response to HF diet are age-sensitive. If the diet is initiated in juvenile mice, males show greater metabolic impairments (31). This sex difference is eliminated if the diet is initiated in adulthood (31), and reversed if initiated in middle-age – with females showing greater metabolic impairment (19, 31). In line with our prior findings, in the current study, the diet was initiated in young adulthood and thus WT males and females showed similar levels of metabolic impairment. However, AD/MxD females developed greater metabolic impairments than males in response to HF diet. Contrary to our findings, Barron (2013) found no sex differences in weight gain between male and female 3xTg-AD mice on a HF diet, though males on a HF diet had increased insulin levels while females did not (21). This enhanced metabolic effect observed in AD/MxD females in the current study is reminiscent of our findings in middle-aged mice in which females are more adversely affected than males. One possibility is that the AD and MxD females might have displayed a form of accelerated aging of their metabolic system and/or brain regulation of metabolic function. We have previously reported this female-specific interaction between the effects of a HF diet and the 3xTg-AD genotype that worsens metabolic impairment and examined potential underlying mechanisms (29). In a subset of mice from the current study (WT and AD groups with the sham surgery), we reported markedly increased hypothalamic expression of GFAP and IL-1β, as well as GFAP labeling in several hypothalamic nuclei that regulate energy balance in the HF-fed 3xTg-AD females (29). Thus, neuroinflammation in the hypothalamus likely contributes to the observed sex difference in metabolic function in 3xTg-AD mice. A limitation in our interpretation of these data is in the generalizability of using a diet composed of 60% fat from lard (saturated fat). Diets of different fat composition or combinations with sugar (a Western diet) may have different consequences. Overall, our results show that a chronic 60% fat diet caused greater metabolic impairment in AD/MxD females compared to males.

### Cognitive findings

Although HF diet elicited some cognitive deficits in both sexes, HF-fed females displayed a wider array of cognitive impairments, even in WT mice. While numerous studies have shown that HF diet exacerbates cognitive impairment in AD (20–23, 25), few have examined sex differences. Here, we report that HF diet caused sex-specific cognitive deficits in spatial learning. AD males, regardless of diet, did not show impairment in spatial learning; however, AD females showed an impairment in spatial learning only when fed a HF diet. In both males and females, additive effects of HF diet were observed in spatial memory. In males, LF-fed AD mice did not show spatial memory deficits; however, with a combination of HF diet and AD these impairments emerged. In females, even WT mice showed spatial memory impairment in response to HF diet (as did AD mice). Overall, we found that poorer metabolic measures were also more consistently associated with poorer cognitive function in females, compared to males. In addition to the sex differences observed with HF diet, we also observed a wider array of cognitive impairment in LF-fed AD females compared to LF-fed males. In support of the current data, prior studies have demonstrated exacerbation of cognitive deficits in 3xTg-AD mice by a HF diet (20–25) and greater cognitive impairments in AD females compared to males (34–39). However, there have been some reports greater deficits in in males specifically in regards to working memory (40) and fear conditioning (41). Greater female susceptibility to negative cognitive effects of a HF diet is in line with our prior work demonstrating that HF diet also causes a wider range of cognitive deficits in WT middle-aged females (19). Taken together, our data suggests that females may be more adversely cognitively affected by metabolic disease which could contribute to cognitive deficits during normal aging or dementia.

In both sexes, MxD elicited a wider array of cognitive deficits beyond what was observed in the AD model, with females being most impacted. Among LF-fed mice, spatial learning deficits emerged in MxD females that were not observed in the AD females or MxD males. Among LF-fed males, spatial memory deficits emerged with MxD that were not observed in AD males but were observed in AD/MxD females. Due to floor effects, we were unable to detect effects of MxD in females for some tests or to detect the effect of HF diet in the MxD groups in either sex for most tests. However, it is important to note that MxD females were impaired in all four cognitive tests, while MxD males had preserved spatial learning and ability to perform activities of daily living. Our MxD model involved a unilateral common carotid artery occlusion surgery, which leads to chronic cerebral hypoperfusion (42–44) and cognitive deficits even in WT mice (19, 42, 45, 46). While others have examined the pathological effects of chronic cerebral hypoperfusion in 3xTg-AD mice (47), we are the first to examine sex differences or cognitive effects in this model. Our data support that added vascular pathology in MxD further exacerbates cognitive deficits, compared to AD alone, and that females may be more adversely cognitively affected by chronic cerebral hypoperfusion in MxD.

### Neuropathology findings

In assessing underlying neuropathology, we found that astrogliosis and Aβ accumulation were more severe in AD/MxD females, compared to males. First, we assessed markers of neuroinflammation. Female AD/MxD mice had greater astrogliosis than males in several regions of the hippocampus. We, and others, have previously reported increased astrogliosis in 3xTg-AD mice (29, 34) and in cerebral hypoperfusion mouse models of VCID (46, 48, 49). In comparison to our 7-month-old mice, greater microglia activation has been reported in 12-month-old 3xTg-AD females compared to males (50), thus sex differences may have emerged at later timepoints. Although there were no sex differences in the extent of microgliosis, we did observe sex differences in the relationship between microgliosis and metabolic markers. In females, but not males, more severe glucose intolerance was associated with more severe microgliosis in the CA1 region of the hippocampus. In addition to neuroinflammation, we assessed other classic AD neuropathology, including Aβ and phosphorylated tau. While we did not observe sex differences in phosphorylated tau, we did find that cortical Aβ was increased in AD/MxD females compared to males. Further, MxD (4 months of chronic cerebral hypoperfusion) increased Aβ beyond what was observed in AD mice. Others have found no hypoperfusion-induced elevation in Aβ levels in 3xTg-AD mice (47), however this was after only 2 months of hypoperfusion suggesting that changes we observed may have emerged at a later timepoint. While we did not observe an effect of diet on Aβ levels, others have found that HF diet increases Aβ in 3xTg-AD mice (21, 24, 25, 51), with some studies showing that this effect is specific to females (52, 53). We are limited in our interpretation because we cannot confirm that these cells are neurons and the presence of Aβ in different cell types may have different consequences for pathology. Due to the age of the animals, Aβ plaques were not observed. Intraneuronal Aβ has been observed in 3xTg-AD mice beginning at 4 months of age (54, 55) and frequently appears in neurons that contain neurofibrillary tangles (56). Others have found increased Aβ levels in female 3xTg-AD mice (21, 34, 50, 52, 53, 57) as well as increased Aβ positive cell counts in the dorsal hippocampus (38). While further sex differences in dementia pathology may develop with age, increased Aβ and astrogliosis in females supports the heightened sensitivity of females to AD/MxD and may underlie the wider array of cognitive deficits in females. (52, 53)

In addition to AD pathology, we also assessed the degree of chronic cerebral hypoperfusion. We found that unilateral carotid artery occlusion surgery (MxD mice) caused deficits in cerebral blood flow in the cortical surface that persisted 4 months after surgery. This is in line with our previous work, and that of others, documenting prolonged deficits in blood flow in the ipsilateral hemisphere (19, 42–44, 58). While we observed the expected effect of MxD to cause hypoperfusion, we did not observe any sex or diet differences. However, greater metabolic impairment was associated with more severe hypoperfusion in females, but not males. Blood flow recovery following unilateral carotid artery occlusion surgery could be affected by the degree of collateral blood flow, angiogenesis, and compensation following surgery. A limitation of the current study is that we only examined blood flow in the cortical surface; there may be differences in blood flow in deeper brain structures that we were unable to detect. Vascular dysfunction is both a contributor to and a result of AD pathology. Vascular abnormalities observed in humans with AD and in the 3xTg-AD mouse, such as reduced cerebrovascular volume (59), may make 3xTg-AD mice more susceptible to vascular insult. While we did not directly examine the mechanistic link between AD and vascular pathology, mechanistic possibilities include hypoxia-induced increases in BACE1 expression via Hif1-alpha (60, 61), blood brain barrier breakdown (62, 63), inflammation (43, 64), or altered glucose uptake into the brain (65). Sex differences in the 3xTg-AD mouse have been noted, including a greater association between plaques and markers of hypoxia (50). These findings suggest that there could be a stronger connection between vascular dysfunction and Aβ pathology in females. Our finding that metabolic impairment is associated with more severe hypoperfusion in females only, prompts further research into the potential of metabolic disease to interfere with blood flow compensation and recovery in a sex-specific manner.

There may be additional neuropathologies not addressed in the current study that could have contributed to sex differences in cognitive function. For example, we have previously shown that HF diet impairs adult hippocampal neurogenesis in WT females, but not males (18). This deficit occurred in the dorsal hippocampus, where neurogenesis supports spatial learning and memory. Additionally, HF diet can accelerate AD pathology in 3xTg-AD mice through several mechanisms not assessed in the current study, such as by increasing neuronal cell death, oxidative stress (23), and brain atrophy (20). Future work is needed to find mechanistic insights behind the sex differences we and others have observed. Given the greater AD pathology in females, the interconnectedness of vascular and AD pathology, and our current findings showing that female MxD mice are more adversely affected, MxD may pose a greater cognitive threat to women.

### Potential Mechanisms Underlying Observed Sex Differences

Sex hormones may influence dementia on many levels, in both men and women, and their effects must be viewed with consideration to age and disease state. Estrogens are present in both sexes but are higher in reproductive-age women. Estrogen also protects against AD risk factors (diabetes) and VCID risk factors (hypertension, obesity, diabetes, stroke) in women (66). It is unknown how estrogen would impact AD and VCID when the pathologies overlap in MxD. Generally, estrogen can protect the brain through its vasodilatory, anti-apoptotic, antioxidant, and anti-inflammatory actions (66, 67). Rodent studies have also demonstrated the protective effects of estrogen against Aβ (51, 68, 69). Estrogen treatment in a male hypoperfusion model improved cognitive function and decreased AD pathology (70), but the effect in females is unknown. While the evidence for the protective effects of estrogen is robust, we have found that AD/MxD female mice are more strongly impacted by HF diet. The 3xTg-AD genotype may diminish the capacity of the brain to produce estrogen, given that women with AD have lower levels of brain aromatase (the enzyme that produces estrogen) (68, 71). Additionally, women with AD have lower levels of brain estrogen (71). There is evidence that estrogen’s effect may turn from protective to damaging when acting in the presence of inflammation or a disease state. For example, estrogen can increase inflammation if administered in older mice after a long period of estrogen withdrawal (72) or lose effectiveness in ameliorating AD pathology (51). Additionally, studies examining the impact of hormone replacement therapy on dementia risk have flagged diabetic women as a population in which HRT increases dementia risk (73). Diabetes has been shown to diminish and sometimes reverse the protective effect of estrogen rodent models of ischemia (74, 75). In this study, chronic HF diet-induced obesity could have created an “accelerated aging” effect in females that negated the protective influence of estrogen.

Androgens may also contribute to sex differences reported here. In men, age-related androgen decline is seen as a risk factor for developing AD (76). In male 3xTg-AD mice, gonadectomy (which reduces testosterone) increases Aβ and tau pathology but is rescued by treatment with testosterone (78, 79). While the effects of testosterone on MxD specifically is unclear, testosterone has a complicated influence on the cerebral vasculature (67, 80) with effects that may be harmful, such as increasing vasoconstriction (81–83) and inflammation (84, 85), but decreasing inflammation in disease states (86, 87) in males. The effect of testosterone on cerebrovascular damage resulting from middle cerebral artery occlusion (a model of ischemia) is dependent on age; it is protective in older age but damaging if applied at supraphysiological levels in younger male mice (80, 88–90). Conversely, men with low testosterone are at greater risk of developing diabetes (78, 91, 92), metabolic disease (93), and dementia (76, 94). Our lab has previously reviewed the cerebrovascular actions of androgens and how that relates to metabolic and cardiovascular disease (80). More research is needed to determine if testosterone level changes in prediabetes affect long term risk of MxD.

### Clinical Relevance

Given that there are well documented sex differences in both AD and VCID, sex differences likely play a role in MxD. In fact, women are more likely to develop MxD, according to a 2007 report (95). Prediabetes research is clinically important because the prediabetic population is growing, there is accelerating awareness of its connection to dementia, and, importantly, it is a potentially modifiable dementia risk factor. To incorporate this risk factor into clinical prevention and treatment of dementia, more research is needed particularly in its effects on MxD, which is inadequately represented in research given its clinical prevalence. Women with diabetes have greater risk of developing cognitive impairments, and in certain types of dementia experience greater cognitive impairments than men with diabetes (3, 96, 97). Metabolic disease in general appears to be a greater dementia risk factor for women (66, 67). This also extends to prediabetes, which is an all-cause dementia risk factor (99, 100). A 2021 study found that prediabetes is associated with accelerated dementia onset and declining cognitive function only in women (33). A major difference in how diabetes and prediabetes are handled is that diabetes is more aggressively treated; this may manifest in differences in long term consequences. For example, a recent study found a relationship Aβ burden and prediabetes but not diabetes (101). The authors hypothesized that this is due to beneficial effects of diabetes treatment. It is clear that further research is needed to understand how prediabetes fits into the overall picture of sex differences in dementia.

### Conclusion

In summary, using a mouse model of mixed (AD + VCID) dementia, AD and MxD females showed a wider array of cognitive deficits, compared to males. Astrogliosis and Aβ pathology were also more severe in AD/MxD females, compared to males. When challenged with a HF diet, AD or MxD females also had increased metabolic impairment compared to males. Metabolic impairment was also more consistently associated with reductions in cognitive function in females. More severe glucose intolerance was associated with worse microgliosis in females only. Here, we demonstrate the importance of considering how sex modulates the relationship between risk factors and dementia. This work supports the importance of prediabetes as a risk factor for multiple forms of dementia, particularly for women, and emphasizes increased cognitive and pathological sensitivity to high fat diet. Future studies assessing the overlap of other risk factors, particularly at midlife, will be important for the development and utilization of therapeutic strategies for dementia treatment and prevention.

## Acknowledgements

The authors would like to thank Nathan Albert, David Riccio, and Richard Daniel Kelly for their assistance with tissue collection and imaging and Melissa Thomas for feeding and weighing mice.

## Abbreviations

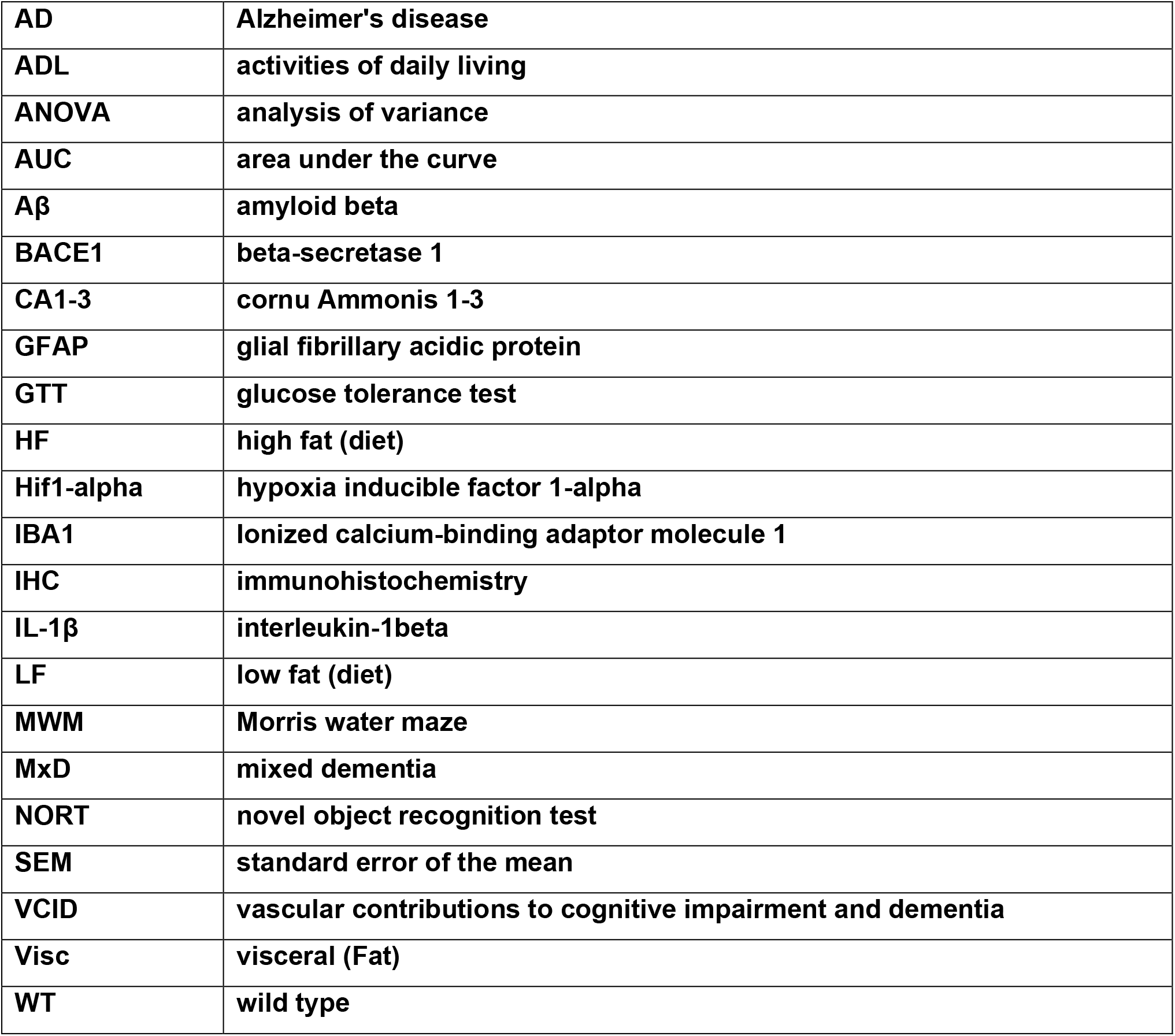

## Authors’ Contributions

KLZ obtained funding for the experiments. LSR, OJG, and KLZ designed the experiments. OJG and AES performed the animal work. OJG, LSR, AES, and CAG performed the experiments. LSR, OJG, CAG, FM, AT, RB, and JO analyzed the data. OJG prepared the figures. OJG prepared the manuscript. LSR and KLZ edited the manuscript. All authors approved the final manuscript.

