## Supplemental Figures for "High fat diet exacerbates cognitive decline in mouse models of Alzheimer’s disease and mixed dementia in a sex-dependent manner"

| Significance Values for 3-way ANOVA | | | | | | | | | | | | |
|---|---|---|---|---|---|---|---|---|---|---|---|---|
| Main effects | Amyloid | Tau | | | GFAP | | | | Iba1 | | | |
| Source of Variation | Amyloid Cell Count | pTau S199 | Total Tau | pTau S199: Total Tau | CA1 - GFAP %Area | CA2 - GFAP %Area | CA3 - GFAP %Area | Dentate - GFAP %Area | CA1 Iba1 %Area | CA2 Iba1 %Area | CA3 Iba1 %Area | Dentate Gyrus Iba1 %Area |
| DEMENTIA | 0.0093 | 0.4398 | 0.5318 | 0.9039 | 0.0934 | 0.0071 | 0.3836 | 0.0164 | 0.0006 | <0.0001 | 0.0049 | 0.1652 |
| SEX | <0.0001 | 0.6151 | 0.0916 | 0.1523 | 0.0419 | 0.3748 | 0.4265 | 0.2376 | 0.0014 | 0.0015 | 0.0031 | 0.0026 |
| DIET | 0.2818 | 0.9178 | 0.6406 | 0.4899 | 0.6879 | 0.1462 | 0.2436 | 0.9323 | 0.1844 | 0.5344 | 0.3871 | 0.2432 |
| DEMENTIA x SEX | 0.3756 | 0.5796 | 0.617 | 0.227 | 0.0469 | 0.0153 | 0.0055 | 0.0298 | 0.0028 | 0.0037 | 0.0194 | 0.138 |
| DEMENTIA x DIET | 0.7817 | 0.2792 | 0.6267 | 0.5395 | 0.2521 | 0.9725 | 0.8159 | 0.3313 | 0.48 | 0.4601 | 0.4522 | 0.5235 |
| SEX x DIET | 0.5955 | 0.5253 | 0.2864 | 0.6048 | 0.2517 | 0.9512 | 0.671 | 0.1333 | 0.5583 | 0.7982 | 0.7385 | 0.7296 |
| DEMENTIA x SEX x DIET | 0.0576 | 0.6561 | 0.6966 | 0.9824 | 0.2568 | 0.1779 | 0.0761 | 0.144 | 0.0799 | 0.1308 | 0.1829 | 0.2272 |

| Significance Values for 3-way ANOVA | | | | | | | | | | | |
|---|---|---|---|---|---|---|---|---|---|---|---|
| Main effects | Metabolic | | | | | Behavior | | | | | |
| Source of Variation | GTT AUC | % change in BW | Visceral Fat (normalized) | Subq Fat (normalized) | Body weight | MWM-Hidden Trial average pathlength | NOR recognition index | Open Field - Distance | Open Field % Center Time | Nest Building Test | MWM-Probe % time in the target quadrant |
| DEMENTIA | <0.0001 | <0.0001 | 0.0028 | <0.0001 | 0.0008 | <0.0001 | 0.0076 | 0.2114 | 0.0001 | 0.0005 | 0.0007 |
| SEX | 0.0022 | <0.0001 | <0.0001 | <0.0001 | 0.4928 | 0.0007 | 0.9651 | <0.0001 | 0.0309 | <0.0001 | 0.0813 |
| DIET | <0.0001 | <0.0001 | <0.0001 | <0.0001 | <0.0001 | 0.451 | 0.4777 | 0.0043 | 0.108 | 0.2575 | 0.005 |
| DEMENTIA x SEX | <0.0001 | <0.0001 | <0.0001 | <0.0001 | <0.0001 | 0.0004 | 0.9642 | 0.0556 | 0.4709 | 0.001 | 0.6486 |
| DEMENTIA x DIET | 0.0088 | <0.0001 | <0.0001 | 0.0555 | <0.0001 | 0.4098 | 0.2241 | 0.0065 | 0.0457 | 0.1142 | 0.1211 |
| SEX x DIET | <0.0001 | <0.0001 | <0.0001 | 0.2369 | <0.0001 | 0.7287 | 0.8007 | 0.0001 | 0.1499 | 0.0247 | 0.3137 |
| DEMENTIA x SEX x DIET | 0.4455 | <0.0001 | 0.0849 | 0.6196 | 0.0535 | 0.0047 | 0.686 | 0.7168 | 0.5071 | 0.8827 | 0.9192 |

**Supplementary Table 1. Statistical test of sex differences.** Results (p values) of 3-way ANOVAs examining main effects of sex, diet, dementia, and interaction effects. Non-significant results are shaded in dark grey, values trending towards significance are shaded with light grey, and significant values are unshaded.

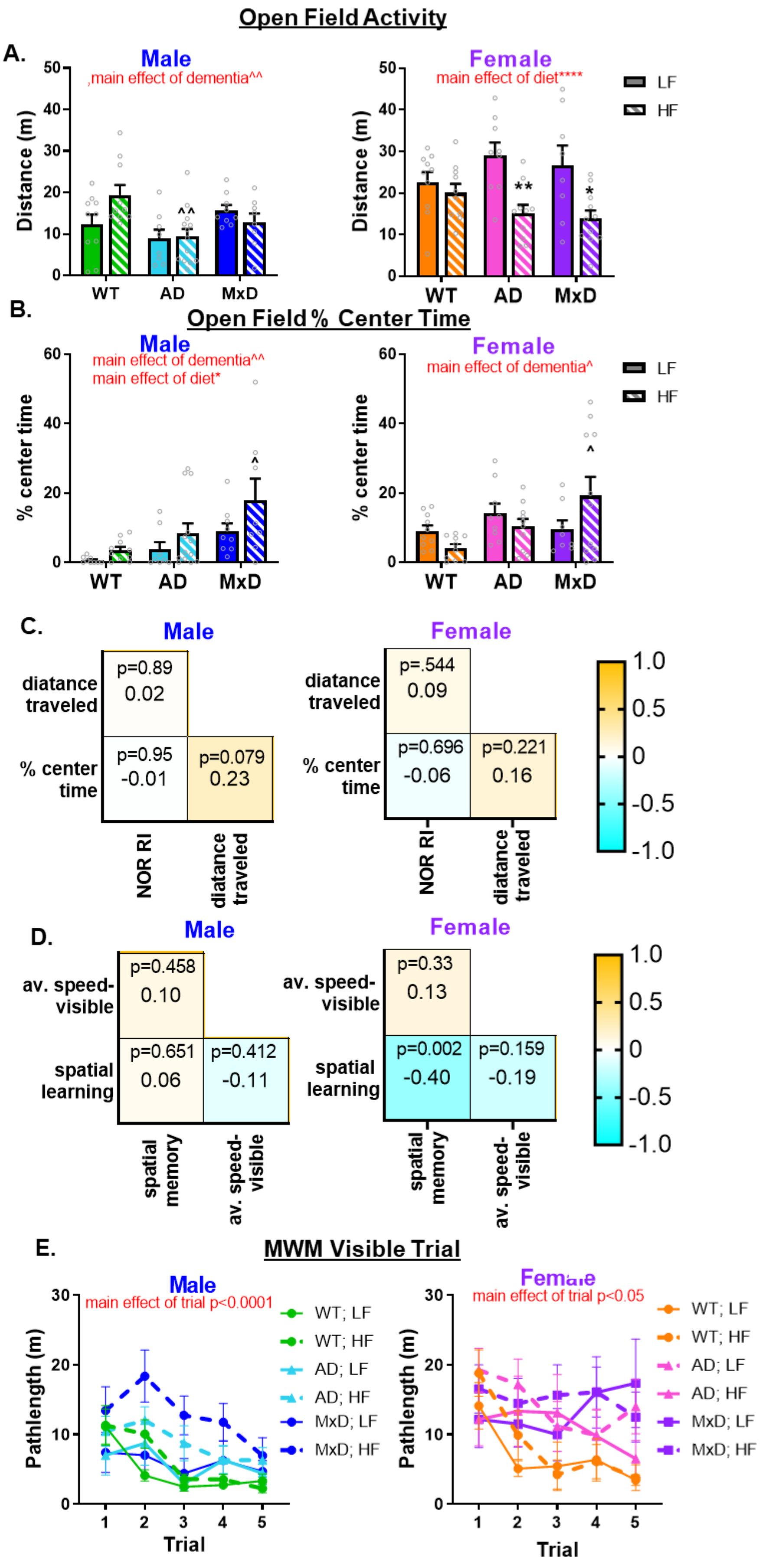
Open Field Activity
A.
Male
,main effect of dementia^^
Distance (m)
WT
AD
MxD
Female
main effect of diet****
LF
HF
B.
Open Field % Center Time
Male
main effect of dementia^^
main effect of diet*
% center time
Female
main effect of dementia^
C.
Male
diatance traveled
% center time
p=0.89
0.02
p=0.95
-0.01
p=0.079
0.23
NOR RI
diatance traveled
Female
p=.544
0.09
p=0.696
-0.06
p=0.221
0.16
1.0
0.5
0
-0.5
-1.0
D.
Male
av. speed-visible
spatial learning
p=0.458
0.10
p=0.651
0.06
p=0.412
-0.11
spatial memory
av. speed-visible
Female
p=0.33
0.13
p=0.002
-0.40
p=0.159
-0.19
E.
MWM Visible Trial
Male
main effect of trial p<0.0001
Pathlength (m)
Trial
WT; LF
WT; HF
AD; LF
AD; HF
MxD; LF
MxD; HF
Female
main effect of trial p<0.05

**Supplementary Figure 1. HF diet decreased locomotor activity in females and increased center time in MxD males and females.** A) General locomotor activity was measured by tracking the distance traveled (in meters) during the open field test. In males (left), there was a main effect of dementia ($p<0.01$, 2-way ANOVA) and in females (right) there was a main effect of diet ($p<0.0001$, 2-way ANOVA). A 3-way ANOVA demonstrated a main sex x diet interaction effect ($p<0.001$, 3-way ANOVA). B) Anxiety-like behavior and disorientation were measured using the %time that the mice spend in the center of the testing arena during the open field test. In males (left), there was a main effect of dementia ($p<0.005$, 2-way ANOVA) and of diet ($p<0.05$, 2-way ANOVA). In females (right), there was a main effect of dementia ($p<0.05$, 2-way ANOVA) and a dementia X diet interaction ($p<0.05$, 2-way ANOVA). C) Correlation matrix for open field measures (distance traveled and % time in the center of the arena) and episodic-like memory as measured by the NOR recognition index (RI) for males and females (n=45-58/sex). No significant correlations between RI and either distance traveled (general locomotor activity) or % center time (anxiety-like behavior) were found in males or females. D) Correlation matrix for average swim speed in the visible trials of the MWM and spatial learning (hidden trial) and spatial memory (probe trial). No significant correlations between average swim speed and either spatial learning or spatial memory in males or females (n=55-57/sex). E. MWM visible trial (day 1) pathlength by trial. There was a main effect of trial (males: $p<0.0001$, females: $p<0.05$, 2-way ANOVA, n=8-11/group). For correlation matrices (C, D): Pearson r values and p values are presented. Yellow: positive correlation, Blue: negative correlation. For A, B, and E: *$p<0.05$ effect of diet, **$p<0.01$ effect of diet, ^$p<0.05$ effect of dementia, ^^$p<0.01$ effect of dementia. Data are presented as mean + SEM (n=8-13 /group).

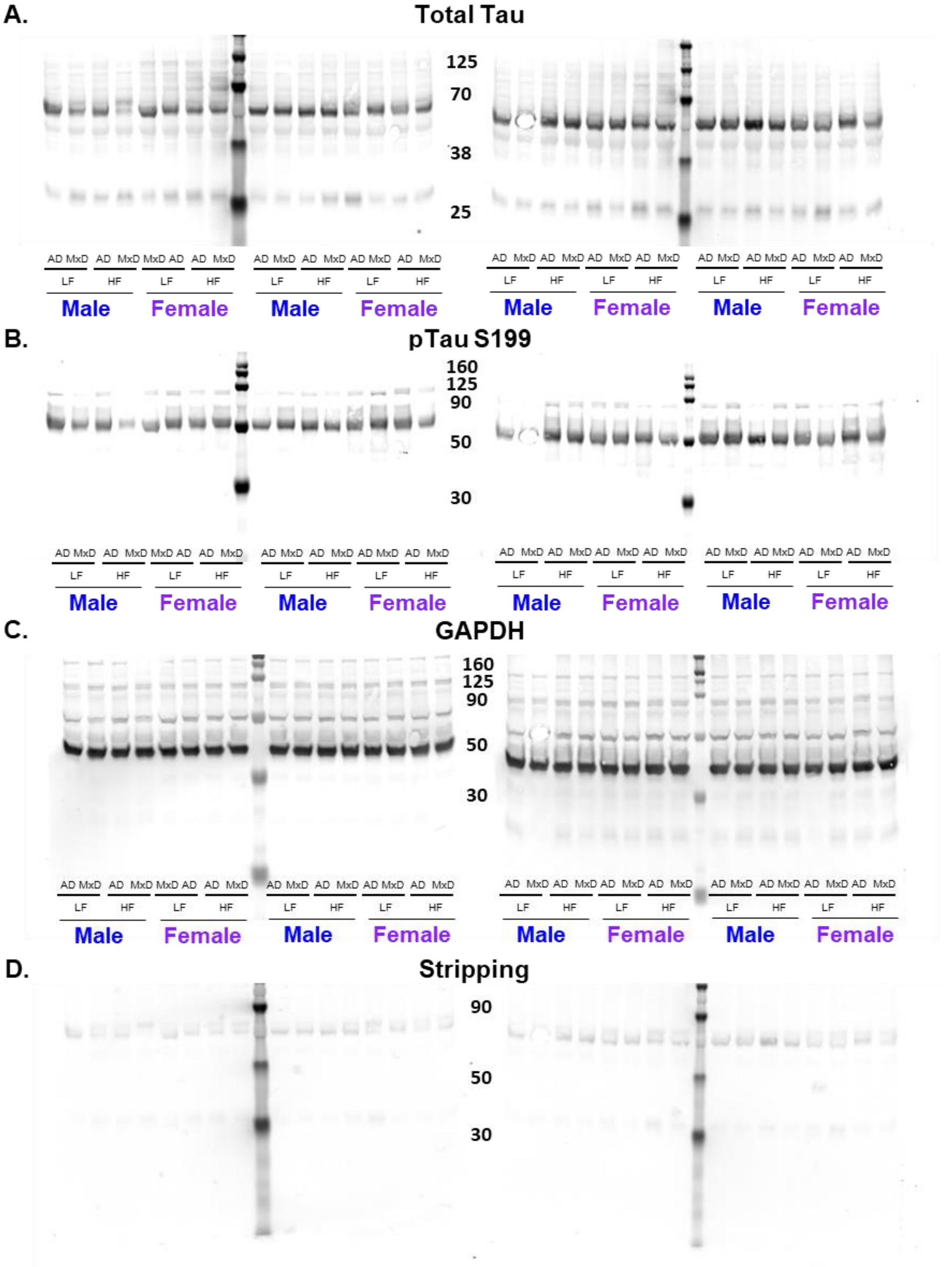


**Supplementary Figure 2.** Western Blot was used to determine the amount of tau protein in right (ischemic for MxD) hippocampal isolates. A) total tau B) pTau S199 C) GAPDH D) stripped gel. n=3-4/group. Images were taken using LI-COR Odyssey CLx.
